## Supplementary figures and images for "Lack of parent-of-origin effects in Nasonia jewel wasp: a replication and extension study"

### S1_Fig.pdf

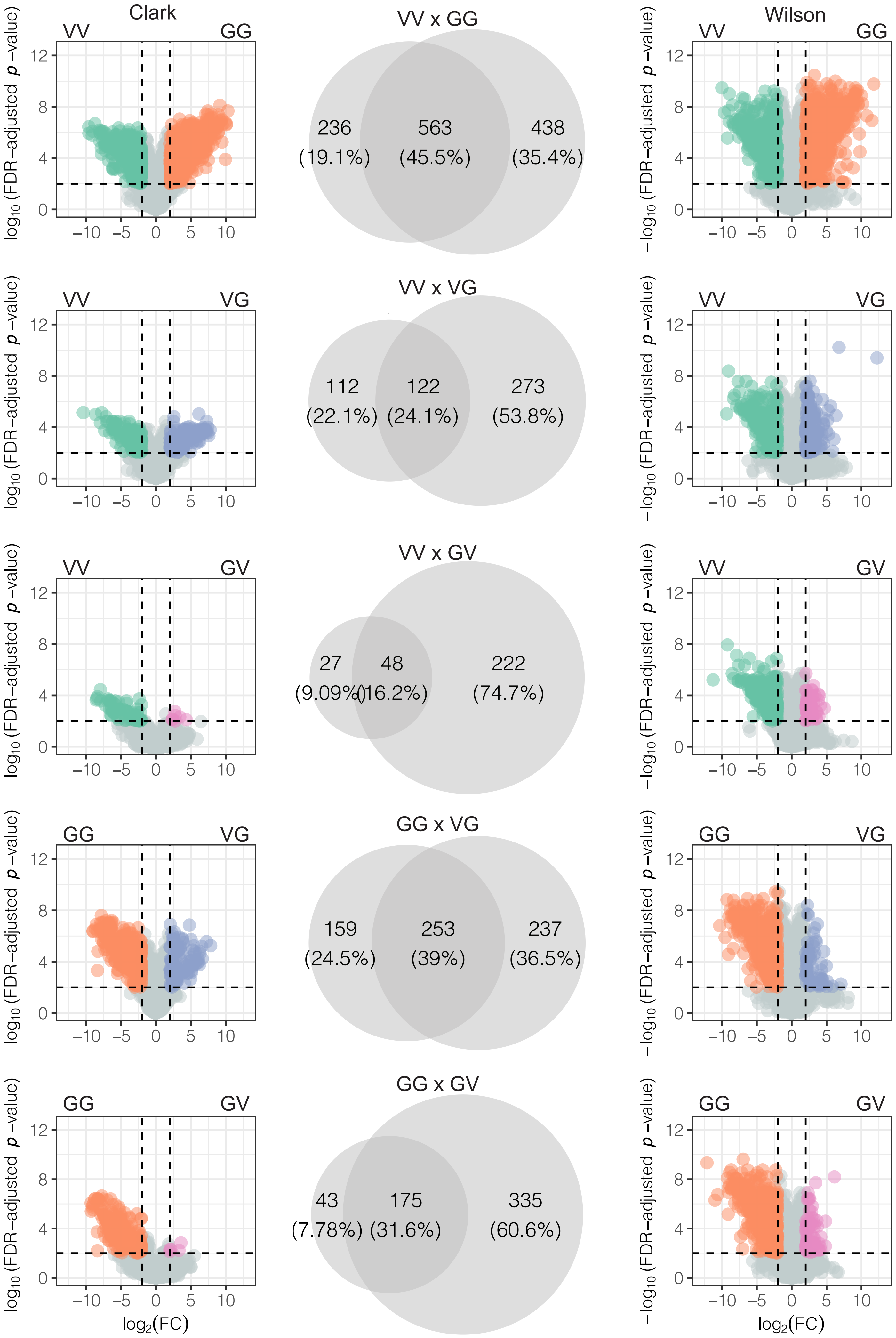
